# Regulation and Function of the HPV16 CircE7 RNA

**DOI:** 10.64898/2026.03.09.710444

**Authors:** Eunice E. Lee, Yihui Huang, Eleanor Dowell, Kaitlyn Walsh, Calvin Hosler, Odera Ifeacho, Doreen Palsgrove, Andrew Day, Richard C. Wang

**Author notes:** Contributed equally (Eunice E. Lee, Yihui Huang, Eleanor Dowell).

## Abstract

High-risk human papillomaviruses (HPV), including HPV16, produce circular RNA that encompass the E7 oncogene (circE7). CircE7 can be detected in HPV16-positive cells and tumors, is preferentially localized to the cytoplasm, is N^6^-methyladenosine (m^6^A) modified and can be translated to produce the E7 oncoprotein. Here, we explored the regulation and function of circE7. Mutation of m^6^A motifs flanking the backsplice junction revealed a single essential m^6^A motif to be essential for circE7 formation. Mutation of this m^6^A motif promoted linear splicing of the E6*I splice site (226^409), suggesting that linear and circular E7 splicing are inversely regulated. Additionally, mutation of an IRES-like motif in circE7 significantly decreased E7 protein expression, without having significant effects on circE7 RNA levels. Knockdown of YTHDC1, but not other m^6^A-binding proteins, decreased both circE7 RNA and protein expression. BaseScope ISH was used to confirm the expression of circE7 in HPV positive head and neck squamous cell carcinoma cell lines. Using both qRT-PCR and BaseScope ISH, we found that serum and amino acid starvation significantly increased circE7 levels. Finally, we generated a HPV16 genome with two point mutations in the circE7 m^6^A motif (Mut2). Stable transduction of primary keratinocytes with Mut2 confirmed the loss of circE7 and increased expression of E6*I. The Mut2 HPV16 genome exhibited significantly decreased viral replication but an increased ability to transform primary keratinocytes. Our studies reveal that the precise regulation of circE7 and E6*I by m^6^A are critical for the ability of HPV16 to infect and transform keratinocytes.

## Introduction

Human papillomavirus (HPV) are highly prevalent oncogenic viruses responsible for up to 5% of all cancers, including up to 80% of oropharyngeal cancers.^1,2^ The oncogenic properties of HPV are largely mediated by the early region of high-risk HPV, especially the E6 and E7 oncogenes.^3^ E6 and E7 work in concert to promote cell-cycle progression through S phase, which enables HPV viral genome replication, while suppressing p53 to block cell cycle arrest and apoptosis. E7 binds to retinoblastoma protein (Rb), inhibiting it and facilitating its degradation, thereby promoting entry into S phase.^4^ The inappropriate entry of cells into S phase can activate cell cycle checkpoints, which stabilizes and increases the levels and activity of the tumor suppressors, including p53.^3^ In high risk HPV, the full-length E6 protein can counter this by binding to p53 to inhibit its transcriptional activity and promote its degradation.^5,6^ However, excess E6 expression has been shown to induce cellular stress responses and inhibit viral replication.^7^ In high risk HPV, alternative splicing of the early region can generate a shorted E6*I isoform, which can inhibit full-length E6 abundance and activity.^8,9^ The regulation of splicing in the early region of HPV is a complex, yet critical, aspect of HPV replication and its ability to transform epithelial cells, whose regulation remains incompletely understood.

Interestingly, some viruses have been reported to generate not only linearly spliced RNA, but also circular RNAs (circRNA).^10–13^ Circular RNAs are produced via backsplicing of a downstream splice donor to an upstream splice acceptor during pre-mRNA splicing.^14^ Their covalently closed structure lacks the 5’ cap or 3’ poly-A tail of linear mRNA and provides circRNAs higher stability than their linear mRNA counterparts. CircRNAs have been shown to perform several biological functions including regulating miRNA, modulating RNA binding proteins, and serving as templates for protein translation.

Previously, we demonstrated that the early region of high-risk HPV, including HPV16, produces a circular RNA, circE7, which contains the E7 open reading frame.^15^ CircE7 is detectable in multiple HPV+ cell lines and primary tumors by transcriptomic sequencing data, RT-qPCR, and Northern blot after RNAse R digestion of linear RNA. Specific knockdown of circE7 decreases cancer cell growth in vitro and in vivo, which can be rescued by exogeneous expression of circE7. Finally, circE7 can be translated into E7 protein based on its association with polysomes in an ATG-dependent manner and decreased E7 protein levels after knockdown of circE7. Notably, circE7 is N^6^-methyladenosine (m^6^A) modified, and this modification is critical to produce circE7 but not linear E7 isoforms.

M^6^A is a common RNA modification that influences stability, splicing, nuclear export, and translation. Several m^6^A-binding proteins have been implicated in the regulation of circRNAs. Specifically, the m^6^A reader, YTHDF3, has been reported to promote translation of some endogenous circRNAs by recruiting eIF4G2 to m^6^A modified circRNAs to initiate cap-independent translation.^16^ YTHDF3 coordinates with other YTH-family m^6^A readers, including YTHDF1, which promote translation, and YTHDF2, which enhances the degradation of m^6^A-RNAs.^17^ Additionally, YTHDC1 is a nuclear m^6^A reader that influences RNA splicing and nuclear export and has been shown to be required for backsplicing of circRNAs.^18–20^ However, the roles of specific m^6^A binding proteins in the regulation of HPV-derived circRNA has not been delineated.

In this study we further characterize circE7. We identify a single m^6^A motif that positively regulates the production of circE7 and inhibits linear E6*I splicing. Translation of circE7 is IRES and YTHDC1 dependent. CircE7 was able to be visualized in HPV16 positive cell lines and primary tissue, and exhibits features of dynamic biological regulation. Transfection of cells with a circE7 deficient HPV genome resulted in increased E6*I RNA level, decreased viral replication, improved cellular immortalization, and elevated levels of p53 protein levels.

## Results

### Discrete m^6^A motif acts as a switch between circE7 and E6*I

In previous work, we demonstrated that m^6^A motifs surrounding the E7 coding region are essential for circE7 formation.^15^ Here, we tested whether a specific m^6^A motif is required for circE7 formation. Given the established role of splice site-adjacent m^6^A motifs in splicing regulation,^21–23,19^ we focused our efforts on the two candidate m^6^A motifs located immediately adjacent to the circE7 backsplice junction. Transversion mutations of the adenosine residue in the m^6^A motifs located near the splice donor (m^6^A-SD-853) and splice acceptor (m^6^A-SA-418) of the circE7 backsplice junction were generated (Fig 1A). Transfection of these mutant circE7 expression constructs revealed that the splice acceptor motif (m^6^A-SA-418) is essential for circE7 RNA production, while the splice donor motif (m^6^A-SD-853) did not show consistent effects on circE7 RNA levels. These effects were reproducible across several different cell lines (Sup Fig 1).

**Figure 1.**
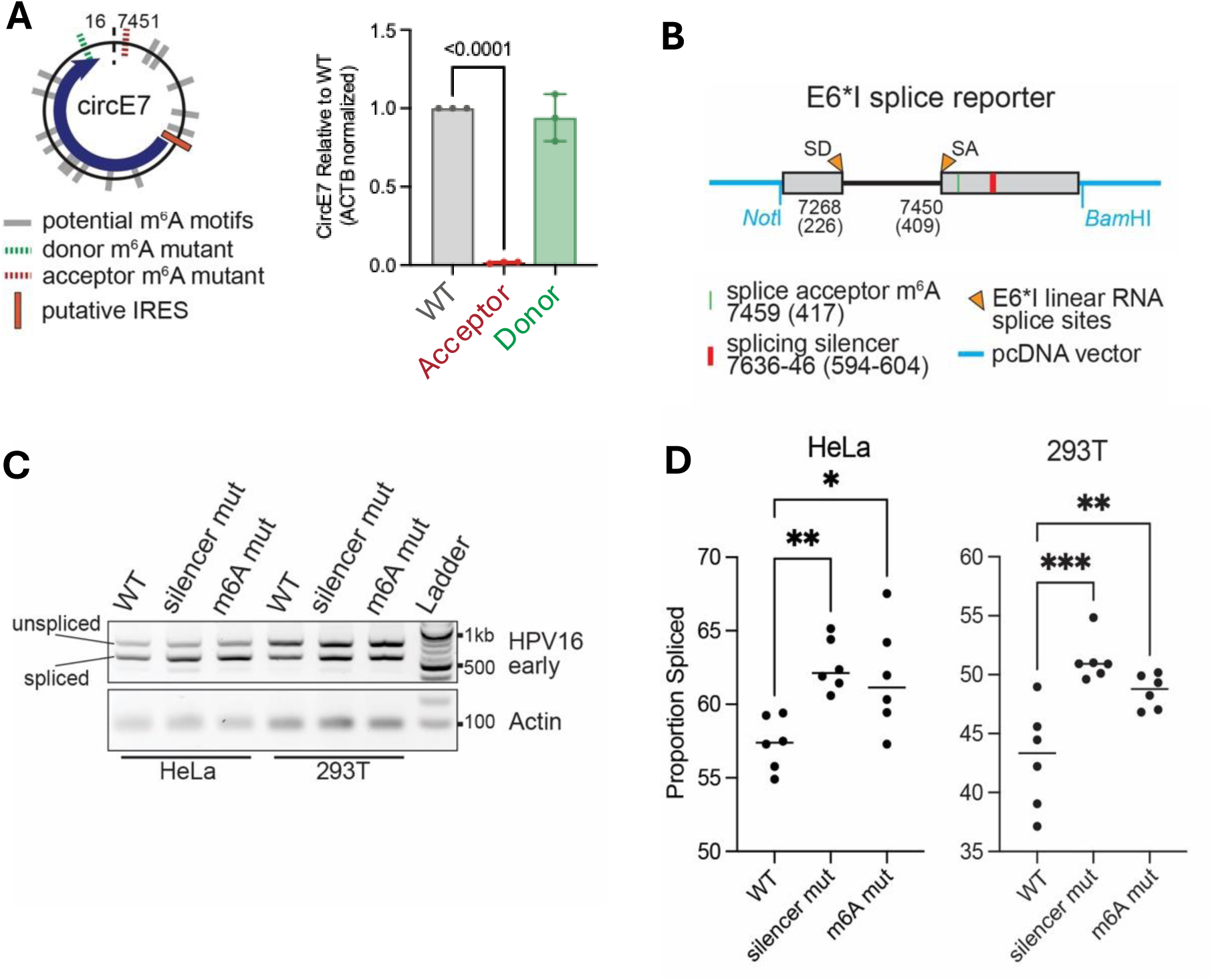
A single m^6^A site coordinates backsplicing of circE7 and linear splicing of E6*I. **(A)** Mutation of the circE7 backsplice acceptor m^6^A (red), but not the backsplice donor m^6^A site (green), significantly decreased circE7 abundance in transfected MDA-1483 cells by RT-qPCR. **(B)** Splicing reporter construct from early region of HPV16 used to assess the effects on splicing of m^6^A circE7 acceptor and E7-silencer mutant (Zheng, J Virol, 2020). **(C)** Endpoint RT-PCR identifies unspliced and spliced early region RNA from cells transfected with either complete (WT), mutated splicing silencer, or mutated m^6^A motif constructs. Both silencer and m^6^A acceptor mutants promote spliced (lower) band compared to the WT sequence. **(D)** quantification of C. *p<0.05, **p<0.01, ***p<0.001, t-test.

To investigate the role of the splicing acceptor m6A motif (m^6^A-SA-418) in linear splicing, we designed a splice reporter plasmid construct containing the previously described E6*I splice site (226^409) but lacking the circE7 backsplice donor (SD880). Two mutant constructs based on this splice reporter plasmid—a splice acceptor m^6^A motif mutant (m^6^A-SA-418) and a previously described splicing silencer mutant—were generated (Fig 1B).^24^ Endpoint RT-PCR from cells transfected with the wild-type reporter construct revealed bands consistent with both spliced and unspliced RNA (Fig 1C-D). As expected, mutation of the splicing silencer motif resulted in increased linear 226^409 splicing. Notably, the single acceptor motif (m^6^A-SA-418) transversion mutant also increased linear 226^409 splicing. These results suggest that the splice acceptor m^6^A motif coordinately suppresses linear 226^409 (E6*I) splicing while promoting circE7 formation.

#### Regulation of circE7 Translation

CircE7 can be translated to produce E7 protein.^15^ After transient transfection, circE7 associates with polysomes, suggesting that overexpressed circE7 is translated. To extend these findings, we assessed whether circE7 associates with polysomes at physiological levels. Polysomes were isolated from two squamous cell cancer cell lines, HPV16-positive SCC154 cells and HPV16-negative MDA1483 cells. RT-PCR of polysome fractions from SCC154 cells detected circE7 RNA, while it was not detected on MDA1483 polysomes (Fig 2A-B).

**Figure 2.**
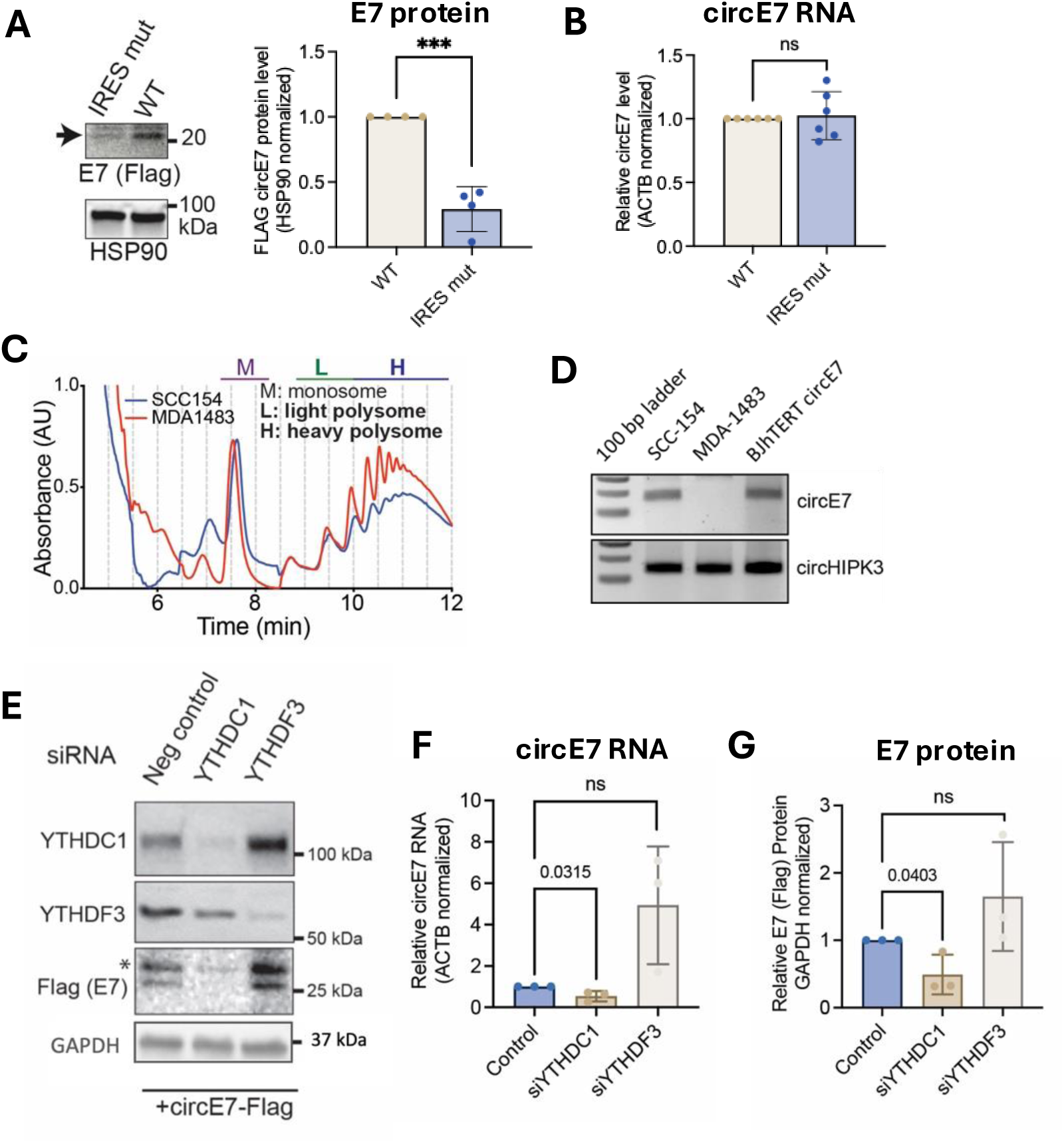
CircE7 RNA translation is dependent on an IRES-like motif and m^6^A binding proteins. **(A)** Western blot from MDA-1483 cells transfected with WT or IRES mut circE7 plasmid. Representative blot (left), quantification (right). ***p<0.001 **(B)** RNA levels of the IRES mutant cricE7 RNA by RT-qPCR in MDA-1483 cells. Difference not statistically significant by t-test. Twice as much RNA was transfected in to get equivalent levels of RNA between IRES and WT. **(C)** Representative tracing of circE7-transfected cells after polysome enrichment assay with the monosome (M), light polysome (L), and heavy polysome (H) fractions indicated. Polysomes isolated from SCC-154 HPV+ (red) and MDA1483 HPV- (blue) cells. **(D)** Endpoint PCR of CircE7 or circHPK3, in light and heavy polysomes from SCC-154 (HPV+) cells, MDA1483 (HPV-) cells, and BJ cells transfected with hTERT circE7. All samples were RNAse R treated. **(E)** Western blot from HeLa cells co-transfected with a circE7-Flag expression vector and the indicated siRNA. HSP90, loading control. *indicates non-specific band. **(F)** RT-qPCR for circE7 RNA levels after YTHDC1 knockdown. **(G)** Quantification of western blot (n=3).

Previous studies revealed that the early (E6-E7) region of HPV16 contains sequences capable of promoting cap-independent translation, possibly through base pairing and interaction with the 18S ribosomal RNA.^25^ Genome comparisons between high-risk and low-risk HPV further narrowed a region complementary to 18S to a short region in the 3’ end of E6, which is contained in circE7.^26^ To test that whether this sequence could function as an IRES-like motif, we generated an IRES-like mutant circE7 in which nucleotides complementary to 18S were substituted with transversion mutations (Fig 2C). WT or IRES-like mutant Flag-tagged circE7 plasmid were transfected into MDA1483 cells. Strikingly, mutation of the IRES-like mutant circE7 markedly reduced levels of E7 protein, despite comparable circE7 RNA abundance (Fig 2D). These results indicate that the primary sequence of circE7 is critical for its translation, possibly through IRES-like elements that mediate ribosome recruitment.

#### The Role of m^6^A Binding Proteins in CircE7 Formation and Translation

Given the critical role of m^6^A sites in circE7 function, we next investigated whether m^6^A-associated RNA binding might also influence circE7 formation. YTHDC1 has been reported to be involved in backsplicing of endogenous circRNA.^20,27^ Co-transfection of cells with the Flag-circE7 plasmid and either YTHDC1 or YTHDF3 siRNA resulted in the expected decrease of YTHDC1 or YTHDF3 protein expression, respectively (Fig 2E). YTHDC1 knockdown significantly decreased circE7 RNA levels (Fig 2F), accompanied by a marked reduction in E7 protein production (Fig 2G). In contrast, YTHDF3, which has been reported to interact with eIF4Gs and promote endogenous circRNA translation,^16^ did not affect cricE7 RNA or E7 protein levels upon knockdown (Fig 2E-G, Sup Fig 2A). Other m^6^A related proteins, including YTHDC2 and hnRNPL, have been implicated in endogenous circRNA formation.^28,29^ However, knockdown of neither YTHDC2 nor hnRNPL did inhibit circE7 formation (Sup Fig 2B-D). Surprisingly, knockdown of hnRNPL increased both circE7 RNA and E7 protein levels (Sup Fig 2B-D), suggesting more complex roles for m^6^A binding proteins in circE7 formation. In summary, we confirm YTHDC1 is the key role in circE7 formation, while other m^6^A -related binding proteins appear to exert more nuanced effects.

#### Detection of CircE7 in vivo and physiologic regulation of CircE7

While NGS and RT-PCR have confirmed that circE7 is expressed in HPV16+ tumors,^15,30,31^ these methodologies do not allow for the visualization of circE7 in situ. To detect circE7 directly in cells in situ, we utilized BaseScope, an RNA in situ method capable of discriminating between RNA at near nucleotide resolution (Sup Fig 3).^32^ Using this approach, we confirmed that distinct E6*I and circE7 isoforms are detectable in the HPV16-positive SCC-154, while neither isoform was detected in HPV16-negative HeLa cells (Fig 3A). Both E6*I and circE7 were also detectable in HNSCC FFPE tissue sections (Fig 3B). In two cases of HPV16-positive HNSCC in which circE7 was identified by sequencing, BaseScope analysis showed RNA ISH specific for both circE7 and E6*I (a-b). In one HPV16-positive HNC tissue section in which circE7 was not identified, E6*I was detected (c). As expected, neither circE7 nor E6*I was detected in the HPV16-negative adnexal tumor control section (d). These tumor-specific differences in the expression of circE7 and linear E6*I suggests complexities in the regulation of E6*I and circE7 splicing.

**Figure 3.**
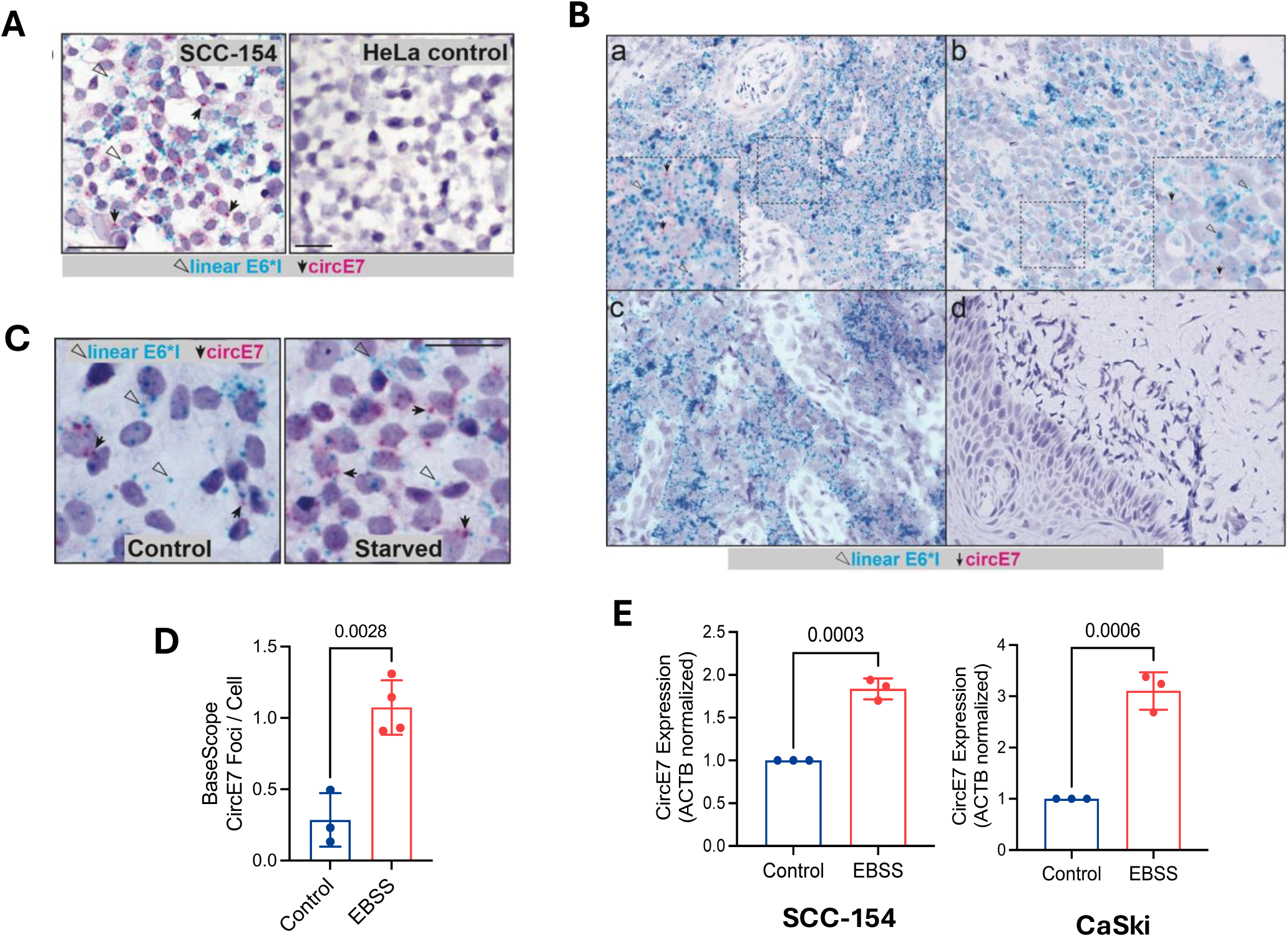
CircE7 RNA is upregulated during cellular starvation. **(A)** SCC-154 (HPV16+) and negative control HeLa cells (HPV16-) were probed by BaseScope Duplex custom probes for either E6*I or circE7. Bar, 50mm. Arrows highlight the presence of circE7 (red) or E6*I (cyan) probes. **(B)** BaseScope detection of circE7 and linear E6*I in HPV+ HNSCC FFPE sections. **(a, b)** Visualization of circE7 (red) and linear E6*I (cyan) in p16+ HNC. **(c)** Linear E6*I (cyan) but minimal circE7 in a p16+ HNC **(d)** No detection of either circE7 or linear E6*I in HPV- adnexal tumor. **(C)** SCC-154 cells were probed for linear (E6*I) or circE7 RNA by BaseScope with or without starvation in EBSS (4 hrs.). **(D)** Quantification of C. Starvation significantly induced circE7 levels as assessed by BaseScope Quantification of BaseScope foci (right) (n=4, unpaired t-test). Bar, 50 mm. **(E)** EBSS starvation induced circE7 levels in SCC-154 (b) and CaSki (c) as assessed by RT-qPCR (n=3, unpaired t-test).

In addition to inter-tumor variability in circE7 expression, even circE7+ tumors showed intra-tumoral heterogeneity of circE7 expression. Tumors exhibit heterogeneous microenvironments that impose distinct adaptive demands on cells located in the core versus the periphery.^33,34^ Previously, we noted that increased cell density augments circE7 expression.^23^ Notably, m^6^A modifications have been reported to increase in response to nutrient starvation.^36,37^ We therefore whether nutrient availability influences circE7 expression by treating SCC154 cells with amino-acid and serum free Earle’s Buffered Salt Solution (EBSS) media.

BaseScope for E6*I and circE7 via showed a marked and significant increase in circE7 levels, with most cells now containing at least one circE7 foci (Fig 3C-D). To orthogonally validate this finding, we performed RT-qPCR with circE7 specific primers and detected a two to three-fold increase in circE7 in starved SCC-154 and CaSki cells (Fig 3E). We conclude that nutrient starvation is a physiological stressor that modulates circE7 levels.

### Effects of CircE7 on HPV16 Genome Replication and Keratinocyte Transformation

To investigate the influence of m^6^A-SA-418 on HPV16 infection and transformation, we generated a mutant HPV16 genome in which m^6^A-SA-418 cannot be methylated. Two substitutions were generated which abolished the m^6^A motif (TAACT to CAATT) without affecting the coding sequence of E6, named Mut2. We transfected wild type and Mut2 genomes into primary human keratinocytes using previously established protocols.^38^ Cells underwent drug selection for six days, after which both WT and Mut2-transduced keratinocytes were able to generate stably populations. Sanger sequencing confirmed successful transduction of the expected wild-type and Mut2 HPV16 genomes.

Both RNA and Hirt DNA extracts were collected from early-passage keratinocytes. Using RT-qPCR, we confirmed that circE7 was markedly reduced in Mut2-transfected keratinocytes (Fig 4A, left). Consistent with our in vitro studies, levels of E6*I increased, further supporting the contention that m^6^A-SA-418 acts as a regulatory switch between circE7 and E6*I (Fig 4A, right). Given the increase in E6*I, we assessed p53 protein levels in Mut2. Western blot revealed that p53 levels were increased in Mut2-infected, consistent with a decrease in E6-mediated degradation of p53 (Fig 4B).

**Figure 4.**
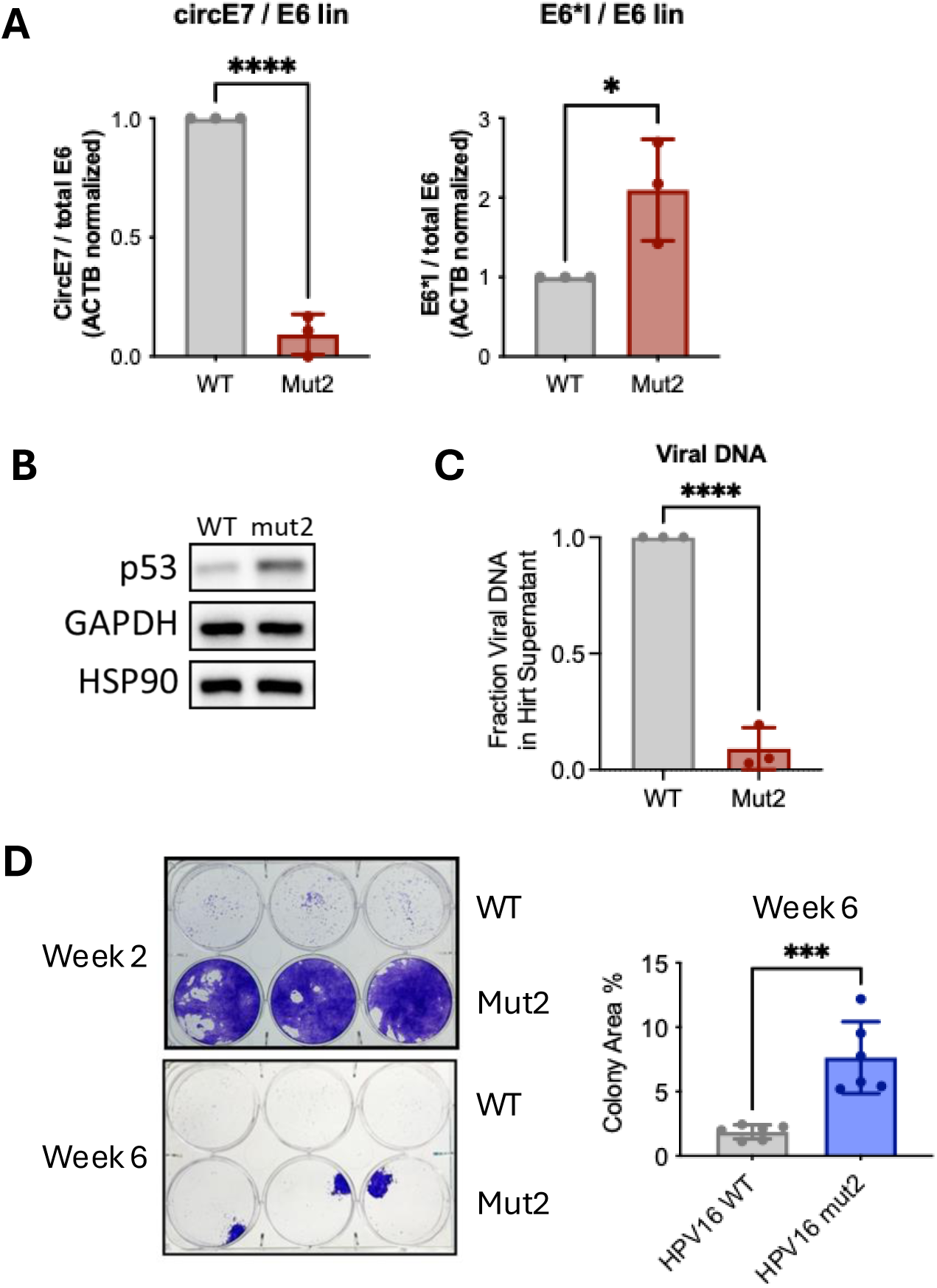
Impact of the circE7 m^6^A motif on HPV16 replication and immortalization. **(A)** RT-qPCR of circE7 and E6*I normalized to linear E6 RNA from primary keratinocytes transfected with HPV16 WT or HPV16 mut2 genomes (circE7 splicing acceptor m^6^A motif mutation, no change to AA sequence). *p<0.05, ****p<0.0001. **(B)** Western blot of p53, GAPDH and HSP90 in whole cell extracts from Keratinocytes transfected with HPV16 WT or mut2 genomes. **(C)** Viral DNA detected by qPCR after HIRT DNA extraction out of primary Keratinocytes transfected with either wild type or mut2 HPV16. ****p<0.0001. **(D)** Crystal Violet staining of keratinocytes grown in 10% BCS DMEM for 2-6 weeks. Cells were transfected with either wild type HPV16 or HPV16 Mut2. Representative images (left) and quantification at week 6 (right). ***p<0.001

We next assessed whether the loss of m^6^A-SA-418 affects viral genome replication. Quantitative PCR of early-passage Hirt extracts revealed that the m^6^A-SA-418 mutant contained approximately 10-fold less viral DNA in Hirt supernatant, (Fig 4C) suggesting significantly impaired viral replication. Finally, we assessed the ability of WT and m^6^A-SA-418 mutant HPV16 to transform primary keratinocytes. Cultures were transferred from basal keratinocyte media to DMEM supplemented with 10% BCS for up to six weeks (Sup Fig 4). Despite impairments in viral replication, the m^6^A-SA-418 mutant HPV showed a significantly increased capacity to immortalize keratinocytes (Fig 4D). After both two and six weeks in DMEM, m^6^A-SA-418 transduced keratinocytes showed superior viability compared with WT HPV16-tranduced cells as assessed by crystal violet staining.

## Discussion

This study provides a detailed model of circE7 regulation and function within the context of HPV infection. We demonstrate that the m^6^A-SA-418 motif acts as a critical regulatory switch for circE7 backsplicing and that an IRES-like motif is required for circE7 translation. Knockdown of the m^6^A reader YTHDC1 decreases both circE7 RNA and E7 protein levels, indicating it likely is an important factor for circE7 backsplicing, while YTHDF3 did not have an impact on circE7 backsplicing or translation. Furthermore, circE7 is physiologically regulated during serum starvation, indicating a dynamic and directed regulation of circE7. Using a splicing reporter, we demonstrate that circE7 backsplicing and E6*I forward splicing are inversely regulated. In the context of the HPV genome, a circE7-backsplicing-defective mutant (Mut2) demonstrates increased relative levels of E6*I transcript and elevated p53, consistent with the established role for E6*I in inhibiting E6 activity. Cells infected with Mut2 showed impaired viral genomic replication but increased immortalization activity. Together, this paints the picture that circE7 is critical for many aspects of HPV function and could act as a critical regulatory node to modulate HPV early region activity in response to cellular conditions.

This study contributes to the growing body of literature that some circRNAs are translated into protein. Previously, we demonstrated that overexpressed circE7 associates with polysomes in an ATG-dependent manner and that shRNA targeting the backsplice junction reduces E7 protein levels, an effect rescued by the exogenous expression of shRNA-resistant circE7.^15^ Here, we build on these observations by demonstrating that circE7 translation is promoted by an IRES-like sequence and that mutation of this sequence significantly reduces E7 protein production. Additionally, we show that circE7 associates with polysomes in HPV16+ cell lines at endogenous levels, supporting that circE7 is translated in native contexts.

In this study, we identified the m^6^A motif, m^6^A-SA-418, which is responsible for regulating circE7 production. While the backsplicing of other circRNA, such as circZNF609, has been shown to be regulated through m^6^A residues.^20^ This is the first study to show circRNA backsplicing to be regulated by a single m^6^A site. Several prior studies have identified a role for the m^6^A reader, YTHDC1, in circRNA formation.^20,27,39^ Our data support a similar role in circE7 biogenesis, although additional m^6^A readers likely regulate its formation. M^6^A modifications have been reported to change dynamically in response to cell stressors, including nutrient starvation.^36^ Similarly, circE7 levels increased in response to starvation and high cell density,^35^ suggesting that circE7, through its m^6^A motif, is dynamically regulated and deployed in response to changing cellular conditions. Our data support a model in which the single m^6^A site, m^6^A-SA-418, acts as a master toggle between circE7 and linear E6*I. Loss of circE7 correlated with an increase in E6*I RNA.

This alternative splicing toggle allows HPV to finely tune E6*I and full-length E6 levels while maintaining E7 protein levels, as E7 can be translated from both E6*I and circE7. Such nuanced regulatory mechanisms are critical for HPV infection, given the compact genome and shared promoters for all early genes.^7^ Our findings complements previous studies on regulators of E6/E7 splicing to provide a more detailed understanding of the regulation of HPV early region transcripts.^38,40,41^ Post-translational regulation of E6 and E7 stability by their interacting partners then adds an additional layer of complexity, which ultimately determines the precise levels of E6 and E7 protein function optimal for viral replication.^42–44^

Several genetic features distinguish low-risk HPV from high-risk HPV in terms of transforming potential, including the differential abilities of E6 and E7 proteins to engage p53 and Rb, respectively, and the presence of a C-terminal PDZ-binding motif in the E6 oncoprotein of high-risk HPV.^45,46^ Interestingly, circE7 RNA isoforms have only been detected in high-risk HPV, raising the possibility that evolution of the high-risk HPV early region and backsplicing are linked. Regardless of its transforming potential, we conclude that regulation of circE7 via m^6^A-SA-418 allows HPV to tune early genes E6 and E7 levels to distinctly modulate individual protein expression from a single pre-mRNA. Additional studies of this densely encoded and tightly regulated genomic region will continue to yield insights on both viral and human RNA biology.

## Materials and methods

### Cell Culture

Primary human keratinocytes were purchased from ATCC (PCS-200-010) and cultured in Dermal Cell Basal Medium (ATCC, PCS-200-030) supplemented with Keratinocyte Growth Kit (PCS-200-040) on Collagen I coated plates. SCC-154 cells (ATCC, CRL-3241) were cultured in DMEM/F-12 GlutaMAX media (Gibco, 10565018) supplemented with 10% FBS, 0.1µg/ml hydrocortisone (Sigma, H0888), and 10ng/ml EGF (Gibco, PHG0311L). HeLa (ATCC, CCL-2), 293T (Takara, 632180), and MDA-1483 (gift from CM Chiang lab) were cultured in DMEM with 10% FBS. CaSki cells (ATCC, CRM-CRL-1550) were grown in RPMI with 10% FBS.

### RNA isolation and cDNA synthesis

RNA was isolated from cells using the RNeasy Mini Kit (Qiagen, 74106) according to the manufacturer’s instructions. RNase R digestion was performed as previously described (Zhao, 2019). For cDNA synthesis, RNA was reverse transcribed using the Superscript IV Synthesis Kit (Invitrogen, 18091050) using the manufacturer’s protocol.

### Endpoint PCR and RT-qPCR

Endpoint PCR performed using the SapphireAmp PCR Master Mix (Takara, RR350B) as previously described (Zhao, 2019). RT-qPCR was performed using PowerUP SYBR Green Master Mix (Applied Biosystems, A25742) according to the manufacturer’s instructions.

### Western blot

For western blot analysis, cells were lysed in Cell Lysis Buffer (Cell Signaling Technologies, 5872S), separated on SDS-PAGE gels, and transferred to PVDF membranes. Membranes were incubated overnight at 4°C with the following primary antibodies: YTHDC1 (Cell Signaling, 77422), YTHDF3 (Cell Signaling, 24206), YTHDC2 (Cell Signaling, 46324), FLAG (Sigma, A8592), GAPDH (Santa Cruz, sc-32233), HSP90 (Cell Signaling, 4877S). Western blots were then incubated with the appropriate HRP-conjugated secondary antibodies (Cell Signaling, 7076S and 7074S) and ECL substrate (Perkin Elmer, 50-904-9323) for signal detection.

### Polysome purification and RNA preparation

Polysome fractionation and analysis were performed as previously described (Zhao, 2019). Briefly, SCC-154 and MDA-1483 cells were grown to 60-80% confluence in 15cm dishes and lysed in polysome lysis buffer (20mM Tris pH 7.4, 5mM MgCl_2_, 100mM NaCl, 0.1% NP-40, 100 µg/ml cycloheximide) in the presence of EDTA-free protease cocktail (Roche, 11836170001). To pellet nuclei, cell lysates were centrifuged at 12,000 *g* for 10 min at 4°C, and the supernatant was transferred onto 10-50% (w/v) sucrose gradient columns and ultracentrifuged at 20,000 *g* for 2h at 4°C. RNA was extracted from collected fractions using TRIzol LS Reagent (Invitrogen, 10296010) according to the manufacturer’s instructions. After isolation, 650ng of RNA was first RNase R digested and then used as a template for random hexamer-primed cDNA synthesis (Invitrogen, 18091050). cDNA products were diluted 1:10 in water for endpoint PCR reactions. **E6*I splice reporter**

A synthetic DNA containing the E6-E7 early region (NC_001526, nt 7140-7900, 16ER_splice_Not_Bam) was cloned into pcDNA3.1(-) as a NotI, BamHI fragment to generate the E6*I splice reporter construct. Subsequently, synthetic DNAs containing transversion mutations in the m^6^A motif (16ER_m^6^A_Not_Age) or a previously described splicing silencer (16ER_silence_mut_Age_Bam) were cloned as NotI, AgeI or AgeI, BamHI fragments, respectively, into the wild-type splice reporter construct. Splice reporter constructs were transfected into the indicated cell lines (293T, HeLa) and total RNA was isolated (Qiagen, mini RNA-easy). Endpoint RT-PCR was conducted to evaluate the E6*I splice region and bands corresponding to the unspliced and spliced RNA were identified. Bands were quantified with ImageJ as a proportion of total RNA.

### IRES mutant plasmid

Primer mediated mutagenesis was conducted on pcDNA3.1(-) circE7_WT and pcDNA3.1(-) circE7_Flag1^15,35^ using the ‘CircE7_IRESmut_F’ and ‘CircE7_IRESmut_R’ primers with the NEB Q5 Site-Directed Mutagenesis kit according to the manufactures’ instructions. Sequences were confirmed by Sanger sequencing.

### siRNA Knockdown

One day prior to transfection, 1 x 10^6^ were plated per well in triplicate in 6-well plates. The following pre-designed siRNA were used: YTHDC1 (SASI_Hs01_00062766), YTHDF3 (SASI_Hs01_00202277), YTHDC2 (SASI_Hs01_00161045), and HNRNPL (SASI_Hs01_00211076). The indicated cells were transfected Lipofectamine 300 with 120 pmol of the indicated siRNA and 2 μg pcDNA3.1(-) Flag-circE7 expression plasmid. After 24 hours, cells were harvested for RNA as previously described. For Western blots, cells were lysed in 200 μL lysis buffer with protease/phosphatase inhibitor (CST). Samples were heated at 80°C for 10 minutes prior to western blot.

### BaseScope and Quantification

The BaseScope^TM^ Reagent Kit (catalog no. 323910; Advanced Cell Diagnostics, Inc., Newark, CA, USA) was used to manually perform the BaseScope^TM^ assay on FFPE tissue sections of MSA and MEC per manufacturer guidelines. Custom RNA probes (Advanced Cell Diagnostics, Inc.) were designed to target fusion transcript-specific exon junctions of interest (BaseScope™ Probes, BA-V-HPV16-circE7, catalog no. 1195271-C1; BA-V-HPV16-E6I, catalog no. 725879-C2). Unstained FFPE 5-micron thick tissue sections were baked in a HybEZ^TM^ (Advanced Cell Diagnostics, Inc.) oven at 60°C for 1 hour, followed by a series of deparaffinization steps. The sections were then left to dry at 60°C for 5 minutes. Once dry, RNAscope® Hydrogen Peroxide (Advanced Cell Diagnostics, Inc.) was applied to each slide at room temperature and left to incubate for 10 minutes. The slides were then washed in distilled water twice. This step was followed by manual target retrieval in which 700 mL of 1x RNAscope® Target Retrieval Reagent was poured into a beaker on a hot plate. Once the solution was warmed to 100°C, the slides were left to incubate in solution for 15 minutes. The slides were quickly transferred into distilled water, washed in 100% ethanol, and left out to dry at room temperature. A hydrophobic barrier was then drawn around the tissue of interest using an ImmEdge^TM^ hydrophobic barrier PAP pen (H-4000). The slides were placed inside the EZ-Batch^TM^ Slide Rack (Advanced Cell Diagnostics, Inc.), where RNAscope® Protease III was added to each sample. The slide rack was then inserted in the HybEZ^TM^ humidity control tray and incubated in the HybEZ^TM^ oven (40°C, 30 min), followed by one wash in distilled water. The MEC and MSA probes were placed in the oven in conjunction with the slide rack to equilibrate prior to use. Excess liquid was then removed from the slides post-incubation, probe mix was added and hybridized (oven at 40°C, 2 hrs.), and slides were washed (1x RNAscope® Wash Buffer twice, 2 min) and stored overnight (room temperature, 5x saline sodium citrate (SSC) buffer). The following day, a series of 8 amplification steps was performed per manufacturer guidelines. RED working solution (1:60, BaseScope^TM^ Fast Red-B to Fast Red-A) was applied (incubation for 10 min, room temperature) and slides were counterstained using 50% Gill’s Hematoxylin I staining solution (catalog no. HXGHE1PT; American MasterTech Scientific) (room temperature, 2 min). Samples were then moved to a staining dish containing tap water (repeated until the slides were clear and sections remained purple), dipped in 0.02% ammonia water (221228; Sigma-Aldrich), washed with tap water, left out to dry (60°C, 15 min), and mounted using VectaMount (64742-48-9; Vector Laboratories). Technical controls were performed on FFPE cultured cell pellets of human HeLa cells using a human-specific housekeeping gene positive control probe a non-specific bacterial (BaseScope™ Probes - BA-Hs-PPIB-1zz - Homo sapiens peptidylprolyl isomerase B (cyclophilin B) (PPIB) mRNA, catalog no. 710171; BaseScope™ Duplex Negative Control Probe- DapB-1ZZ, catalog no. 700141). Adjacent benign tissue was used as a negative internal control.

### Immortalization assay

A synthetic cDNA gBlock fragment (IDT) containing the mutant m^6^A motif (AgeI_m6A_Mut2) was cloned into the genomic HPV16 plasmid, pHPV16ANsL (gift from CM Chiang lab) using *Age*I restriction sites. Positive clones (called Mut2) were confirmed by Sanger Sequencing. Primary keratinocytes were immortalized by HPV16 as previously described by Jönsson et al, 2023.

Briefly, early passage primary keratinocytes were co-transfected with an equal DNA ratio of either pHPV16ANsL WT or Mut2 and pCAGGS-nlsCre plasmids (gift from CM Chiang lab) using TransIT Keratinocyte Transfection Reagent (MirusBio, MIR 2800) according to the manufacturer’s protocol. Cells were selected with G418 for 6 days and maintained in keratinocyte media until the cultures reached confluence. Cells were then selected in DMEM +10% BCS for 6 weeks to induce terminal differentiation. After six weeks of selection, colonies were stained overnight with 0.1% crystal violet in 50% methanol. Colonies were quantified using the ColonyArea ImageJ plugin.

### HIRT isolation

Cells (>1 x 10^6^) were scraped into PBS and washed twice in PBS before snap freezing. Cells were lysed in 200 µl Hirt lysis buffer (10mM Tris-HCl-pH 8.0, 10mM EDTA, 0.6% SDS-store at RT) with gentle shaking for 10 mins at room temperature. After addition of 50 µl of 5M NaCl, tubes were gently inverted and incubated overnight at 4°C. Tubes were centrifuge at full speed for 30 mins at 4°C. The supernatant was collected and treated with RNase A (0.2 mg/ml) for 1h at 37°C. 1250 µl pf NTB (Qiagen) and mixed and DNA was purified using a spin column (Macherey-Nagel, PCR and Gel purification Kit) and elute in 30 µl of elution buffer. The DNA (∼0.5 µL) was digested with 10 unit (1 µL) of DpnI for 16 h, which was inactivated at 80°C for 20 mins. A standard curve using 10-fold serially diluted purified plasmids was used to establish a standard curve and qPCR was run using the DpnI digested samples.

**Supplemental Table 1.**
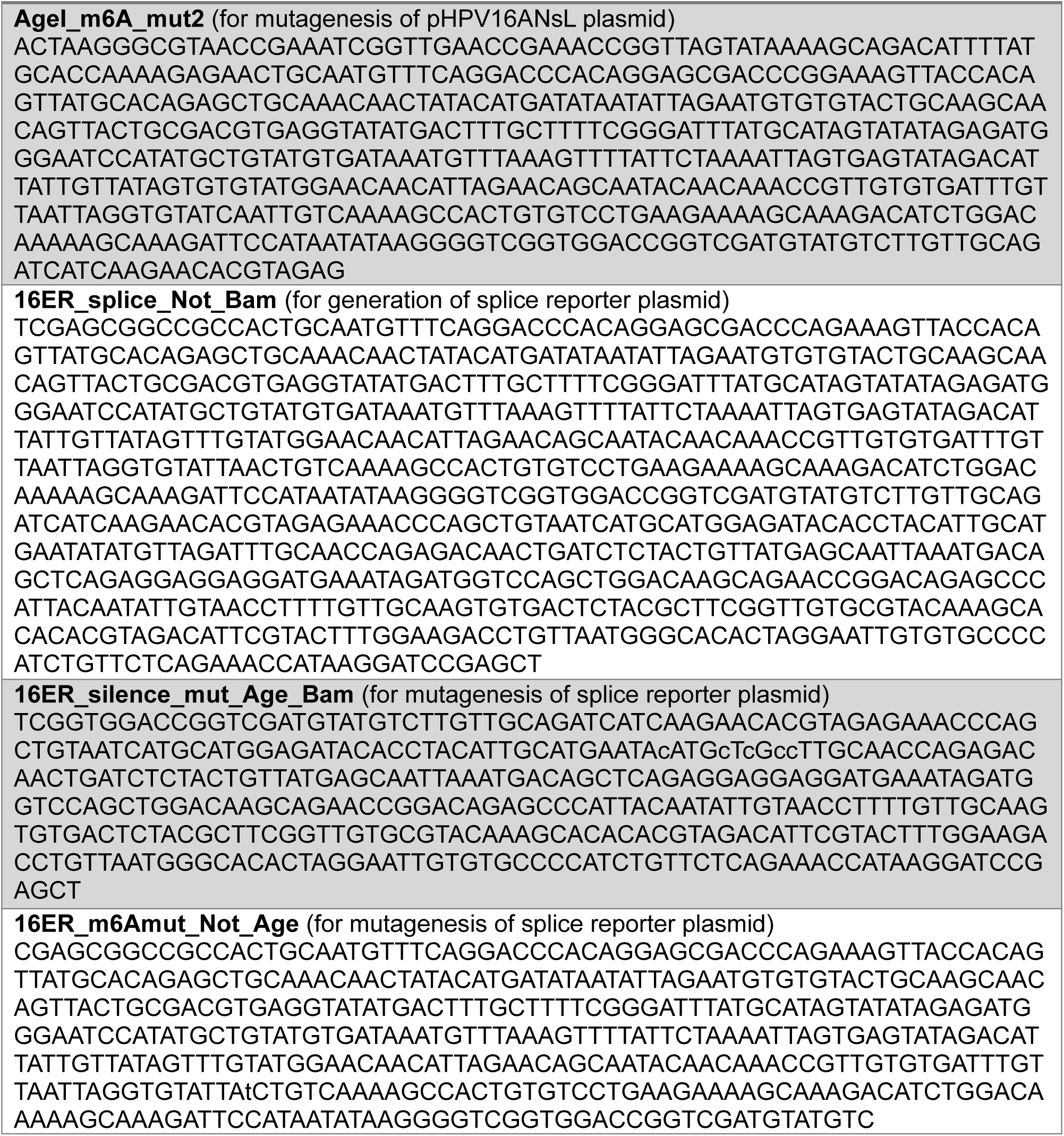
Synthetic DNA sequences.

**Supplemental Table 2.**
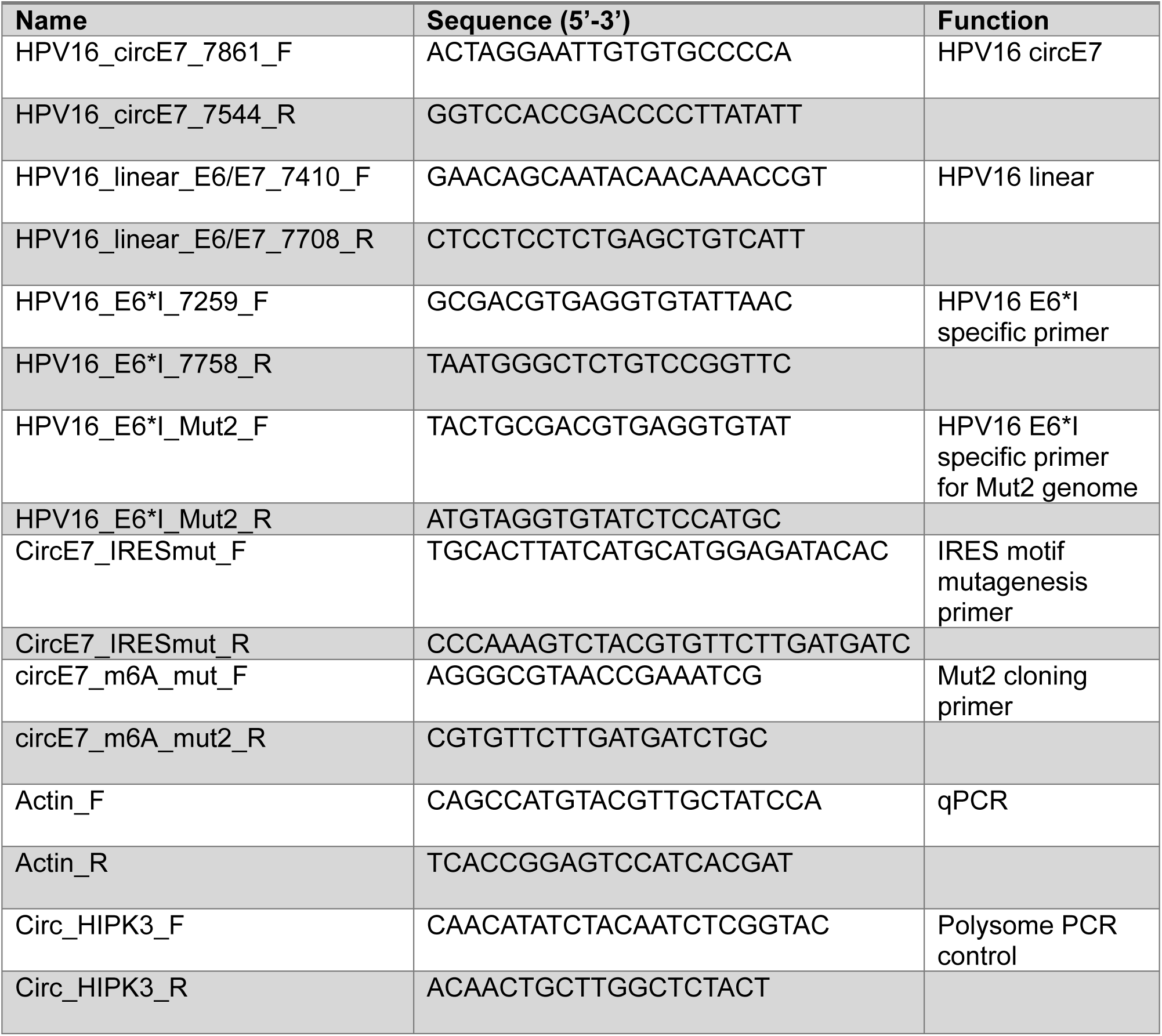
Primer sequences.

---

**Supplemental Figure 1.**
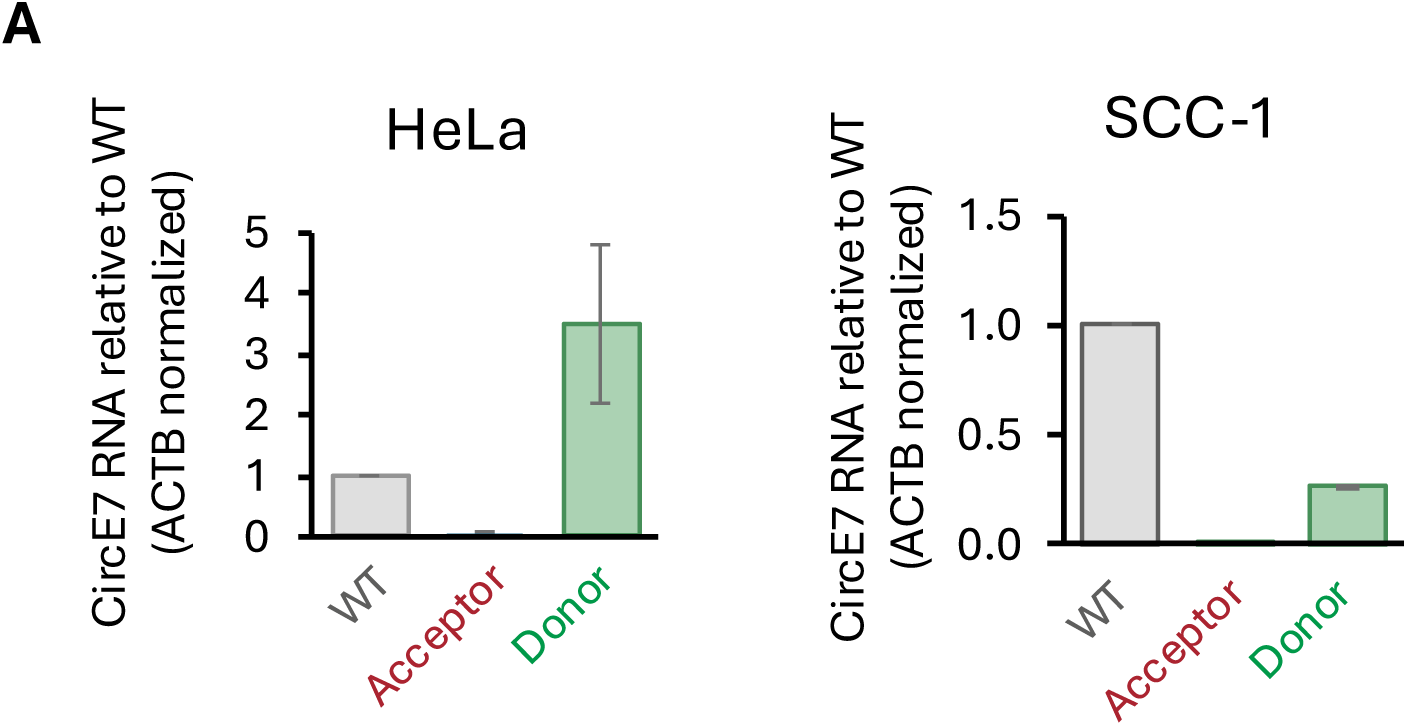
Mutating m^6^A reduces circE7 in multiple cell lines. **(A)** CircE7 RNA assess by RT-qPCR in HeLa and SCC-1 cells transfected with wild type (WT), m6A acceptor mutant (Acceptor), and m6A donor mutant (Donor).

**Supplemental Figure 2.**
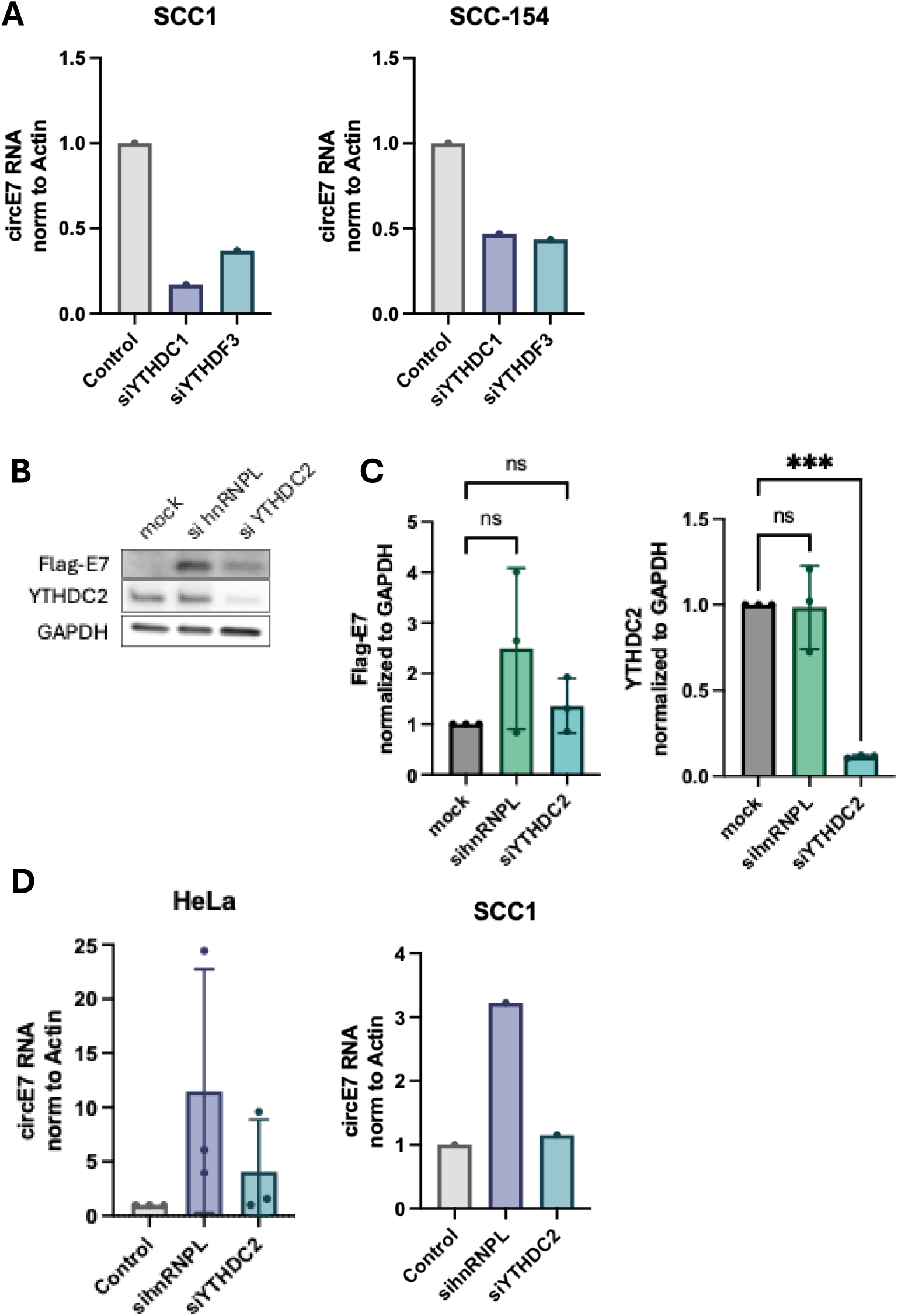
Additional m^6^A readers repress circE7. **(A)** RT-qPCR of circE7 after knockdown of various YTHDC1 or YTHDF3 m^6^A readers in SCC1 and SCC-154 cell lines. n=1. **(B)** Representative western blot of Flag-tagged E7 protein, YTHDC2 and GAPDH proteins after mock, si-hnRNPL, or siYTHDC2 transfection in 293T cells. **(C)** Quantification of Flag-E7 (left) and YTHDC2 (right) western blots. One way ANOVA. ***p<0.001. **(D)** RT-qPCR of circE7 RNA after knockdown of various proteins. Normalized to Actin RNA then control condition. For the HeLa control, si-hnRNPL, and siYTHDC2 n=3; all others n=1. Conditions are not statistically significant.

**Supplemental Figure 3.**
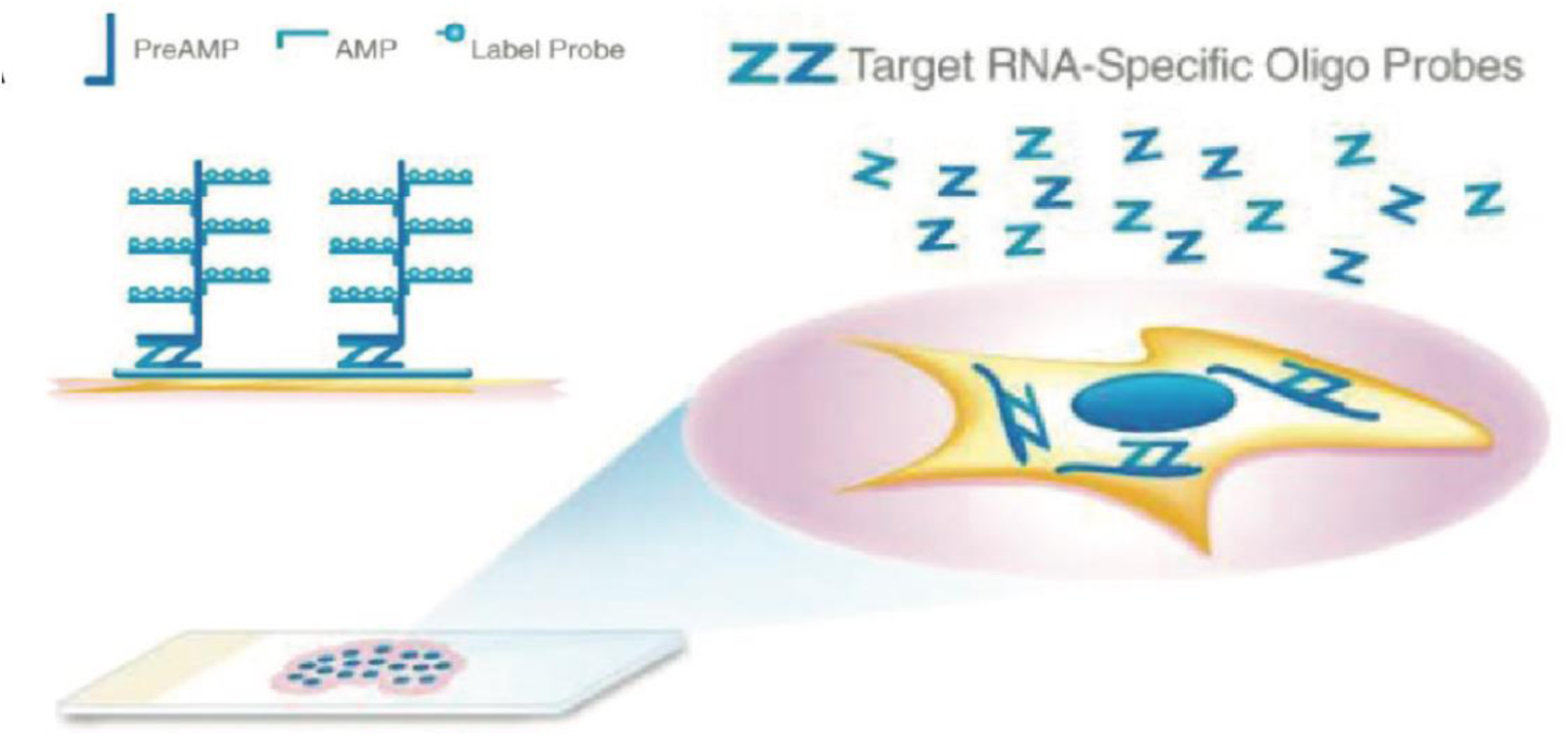
BaseScope schematic. BaseScope RNA-ISH detection of RNA. Double “Z-probes” allow for discrimination of sequences differing by as little as one nucleotide (Abere 2020).

**Supplemental Figure 4.**
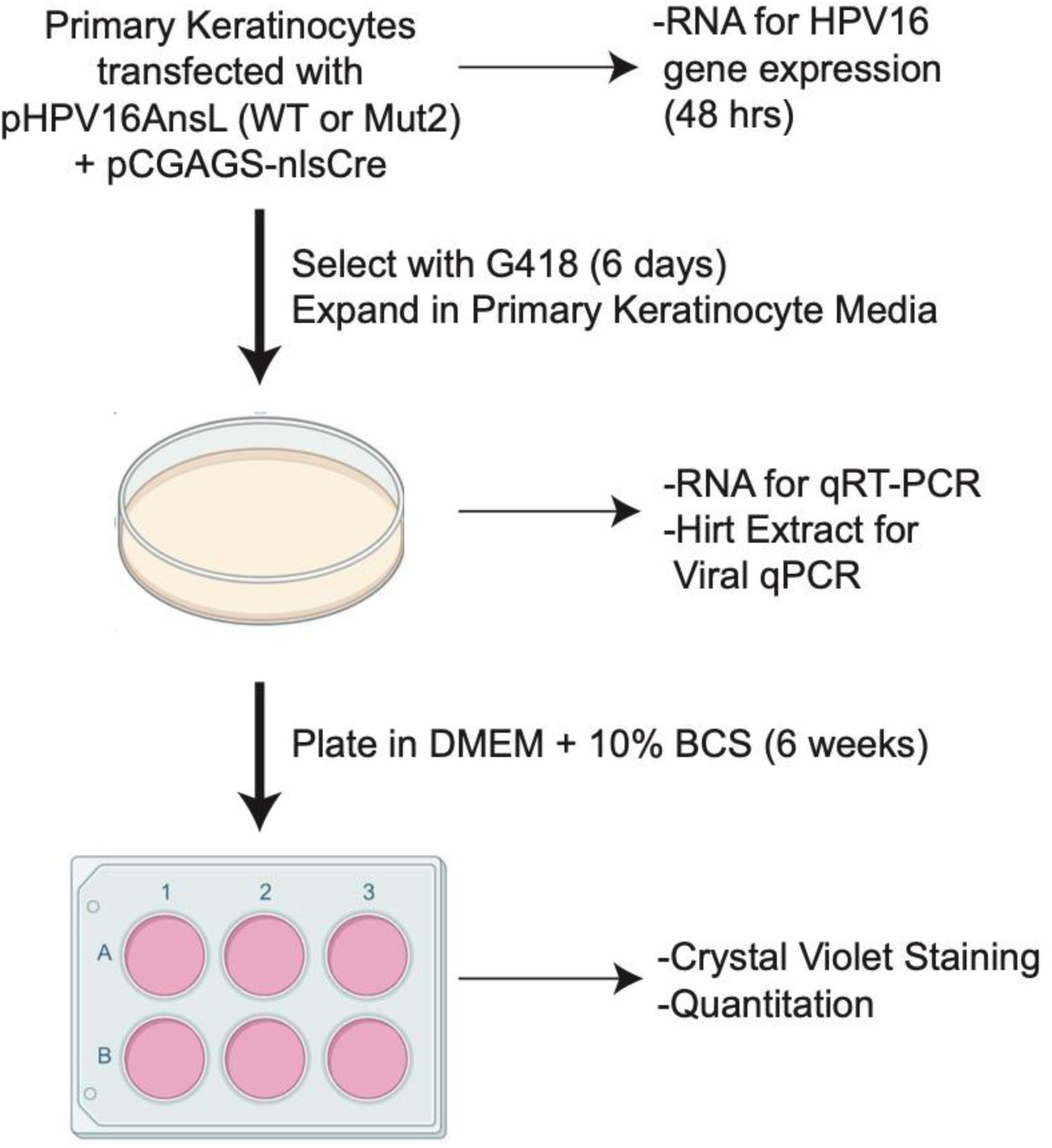
E**x**perimental **design for HPV16 infection evaluation.** Primary keratinocytes were grown in primary keratinocyte media and co-transfected with the pHPV16AnsL genome, either wild type or Mut2, and pCGAGS-nlsCre. 48 hours after transfection, RNA was harvested from transfected keratinocytes for quantifying HPV16 gene expression. Cells were selected with G418 for 6 days and expanded in primary keratinocyte media. After selection, cells were harvested for RNA and Hirt extracts. Selected keratinocytes were then grown in DMEM with 10% bovine calf serum for up to 6 weeks and monitored at various timepoints via crystal violet staining.

